# Novel Probe Design for Multi-Gene Detection Enabled by Cleavage and Capillary Electrophoresis

**DOI:** 10.1101/2025.03.15.643483

**Authors:** Yangqing Gong, Yong Wu, Yong Luo, Xinjiang Lu, Miaomiao Niu

## Abstract

Nucleic acid sequences carry the most fundamental information about life, and gene detection plays a crucial role in studying and applying gene function. Here, we present a novel probe design that utilizes probe cleavage and capillary electrophoresis for gene detection, termed ProbeCE which detects genes by measuring the relative fluorescence signal intensity. Through the evaluation of probes with varying lengths and modifications, ProbeCE is believed to support the simultaneous detection of hundreds of genes. Additionally, the assessment of low-concentration samples has demonstrated the high sensitivity of the ProbeCE method. Compared to the conventional amplification product capillary electrophoresis (AmpCE) method, this new approach significantly increases detection throughput while simplifying primer design and enhancing specificity. In addition, ProbeCE is less expensive than current medium- or high-throughput sequencing for detecting large numbers of known genes.

**GRAPHICAL ABSTRACT:** 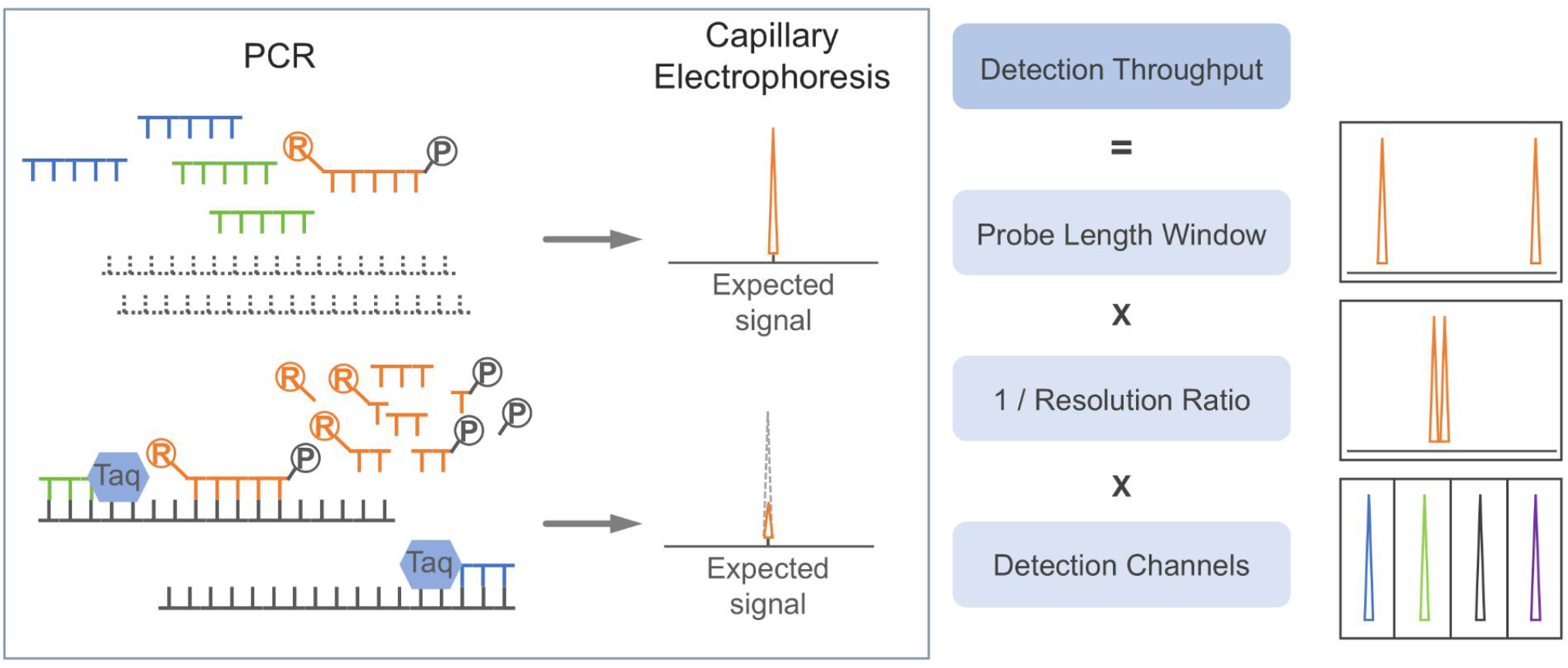

## INTRODUCTION

Multiplex polymerase chain reaction (MPCR) enables simultaneous amplification of multiple DNA fragments by incorporating several primers into a single reaction system. When combined with various detection methods, MPCR enables concurrently identifying many nucleic acid sequences, including the following common methods:

Real-time quantitative PCR (qPCR) consists of two main types: 1. Probe-based qPCR (such as TaqMan, molecular beacons) uses different labeled fluorescent probes to distinguish DNA fragments (1, 2, 3); 2. Dye-based qPCR. The fluorescent dye binds to double-stranded DNA and differentiates nucleic acid fragments based on their melting temperatures (4). When fluorescent-labeled probes are combined with melting temperature analysis, it is theoretically possible to detect up to 30 targets simultaneously (5). However, the melting temperature, which is susceptible to external influences, poses a challenge for the design of densely spaced detection targets, typically limiting most commercial assays to around 20 targets. (6).

Capillary electrophoresis (CE) separates and identifies nucleic acids by leveraging differences in molecular size and charge, which result in variations in migration speed under an electric field (7). While CE can differentiate hundreds of fragments (8), the conventional method, using a combination of PCR and CE is constrained by factors such as unstable fluorescence signal intensity and amplified fragment length, with typical detection capabilities ranging from 10 to 30 fragments (9).

The growing demand for multi-gene detection has driven the development of many novel technologies. Microarray platforms can anchor tens to thousands of probes on a solid surface, allowing for the parallel detection of multiple nucleic acids (10). NanoString nCounter Technology utilizes unique code assigned to each gene under test (11), addressing the limitations of traditional fluorescent PCR, could detect hundreds of genes simultaneously, and enables direct detection of RNA without amplification (12). Next-generation sequencing (NGS) has significant advantages in discovering new genes and conducting functional research and has greatly advanced genome and transcriptome research. Despite the significant reduction in the cost of NGS, its inherent technical requirements demand a multi-step process, which limits the potential for further cost reductions. Furthermore, although NGS is highly effective for whole-genome or transcriptome studies, its capability to target and analyze specific genes is relatively less robust (13, 14).

With the continuous mining of genomic data and the deepening of gene function research, there will be an increasing demand for multi-gene synchronous study and application. Different technologies possess distinct advantages and limitations, and employing a combination of tools can yield more accurate and comprehensive data for a variety of research objectives. Currently, there is a notable absence of convenient and cost-effective methods for the simultaneous detection of tens to hundreds of genes. In this study, we introduce a novel approach designed to address these requirements.

## MATERIALS AND METHODS

### Materials

Standard strains Adv 1 which belong to adenovirus C-type (Adv C), Adv 4 which belong to adenovirus E-type (Adv E), H1N1 (belong to Flu A), Victoria (belong to Flu B), CoV-NL63, Parainfluenza Virus type 2 (PIV 2), Parainfluenza Virus type 3 (PIV 3), Respiratory Syncytial Virus type B (RSV B) were purchased from Guangzhou Standa Biotechnology Co., Ltd (Guangzhou, China). Standard strains of *Bordetella pertussis* (*B. pertussis*) were purchased from the Henan Engineering Research Center of Industrial Microbiology (Henan, China). Plasmid containing specific sequences of CoV-HKU1 was synthesized by Beijing Tsingke Biotech Co., Ltd.

### Nucleic acid extraction and digital quantification

Nucleic acids were extracted using the Magnetic Bead Method Nucleic Acid Extraction Kit (RT-A (SG)-200) from Zhongyuan Huiji Biotechnology Co., Ltd. Digital qPCR was conducted using the qRT-PCR Kit [V7] from Cnpair Biotech Co., Ltd. The reaction mix consisted of 12.5 μL of 2x qRT-PCR Premix, 2.5 μL of 2 μM primers, 2.5 μL of 1 μM probe, 0.5 μL of 25 μM ROX, 2 μL of H_2_O, and 5 μL of template. The PCR protocol was as follows: reverse transcription at 50°C for 10 minutes, pre-denaturation at 95°C for 2 minutes, followed by 45 cycles of denaturation at 95°C for 10 seconds and annealing at 60°C for 60 seconds. This was followed by cooling at 72°C for 5 minutes, an additional denaturation at 95°C for 5 minutes, and signal collection during elongation at 60°C for 1 minute.

### PCR amplification system and procedure using the ProbeCE

PCR reactions were prepared using the DirectFast qRT-PCR Kit (Cnpair Biotech Co., Ltd.). The reaction mixture included 4 μL of DF Reaction Buffer, 2 μL of DF Enzyme Mix, 2 μL of 2 μM primers, 1 μL of probe, 6 μL of H_2_O, and 5 μL of template, for a total volume of 20 μL. Probes modified with FAM or JOE were used at a final concentration of 10 nM, Atto550-modified probes at 20 nM, and Atto590-modified probes at 30 nM. The PCR amplification protocol was as follows: reverse transcription at 50°C for 10 minutes, pre-denaturation at 95°C for 2 minutes, followed by 40 cycles of denaturation at 95°C for 10 seconds and annealing at 60°C for 90 seconds. Primers were shown in **Supplementary Table 1**, and the probes used in this study were shown in **Table 1**.

**Table 1.**
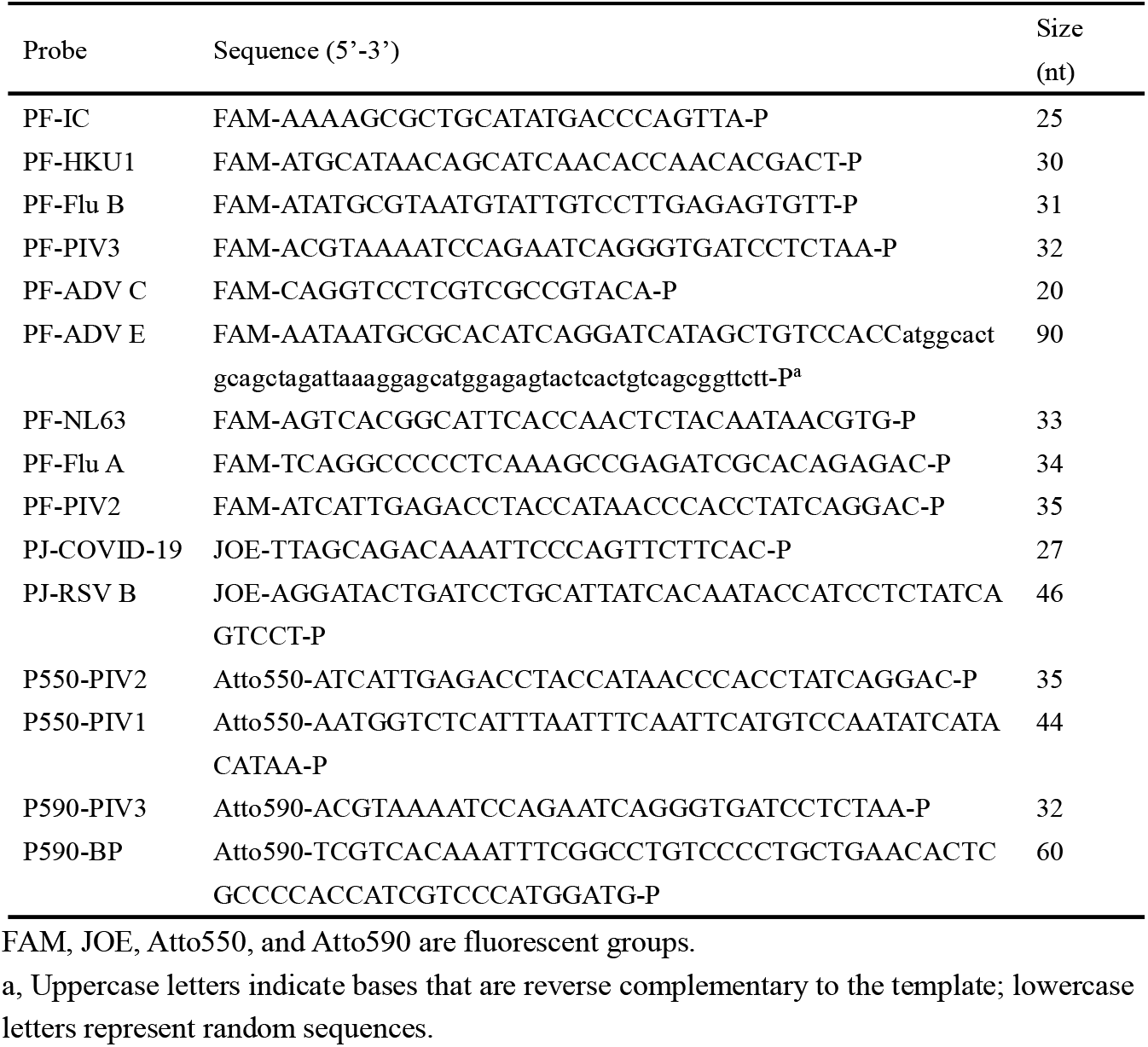
Sequence and modification of probes.

### DNA ladder preparation

Nucleic acid fragments with sizes 20nt, 40nt, and 70nt were synthesized by Invitrogen (Thermo Fisher Scientific Inc.), with 5’-end Alexa Fluor 633 modification and 3’-end phosphate group modification. The sequence of fragments is shown in **Supplementary Table 2**.

### Capillary electrophoresis

Preparation of CE preparation solution: Size 20 (1μM) 5μL, Size 40 (1μM) 5μL, Size 70 (1μM) 5μL, Hi-Di™ Formamide 985 μL. The final CE solution: CE preparation solution 9 μL, PCR amplification product 1 μL. The CE instrument was Applied Biosystems 3500 Dx with the following settings: oven temperature at 60°C, injection voltage at 1.6kV for 8s, prerun voltage at 15kV for 180s, and run voltage at 19.5kV for 1200s.

### Statistical analyses

Three different sample concentration gradient tests were conducted with six replicates each. The data were expressed as mean ± standard error of the mean (S.E.M.) and analyzed using one-way ANOVA. A *P-value* < 0.01 was considered statistically significant. For the probe length range test, also performed in six replicates, the data were presented as mean ± S.E.M. and analyzed using Welch’s t-test. In the signal position stability test, which included six replicates, the results were reported as mean ± range. The multi-channel test was carried out with three replicates. CE raw data were exported using GeneMapper 5.0 software with Amplified Fragment Length Polymorphism (AFLP) settings and subsequently analyzed using GraphPad Prism 7.0.

## RESULTS

### Probe design

Conventional amplification product capillary electrophoresis (AmpCE) designs different-sized amplification windows for each gene and separates amplification products with fluorescent modifications by applying an external electric field **(Figure 1a)**. The novel method ProbeCE uses fluorescently modified probes instead of primers to provide fluorescent signals, and detects genes by measuring changes in signal intensity. The structure of the probe is the key to this method. The 5’-end of the probe is modified with fluorescent groups (can be cut in time), and the 3’-end of the probe is modified with phosphate groups (prevents probe extension and makes the electrophoretic migration position predictable) **(Figure 1b)**.

**Figure 1.**
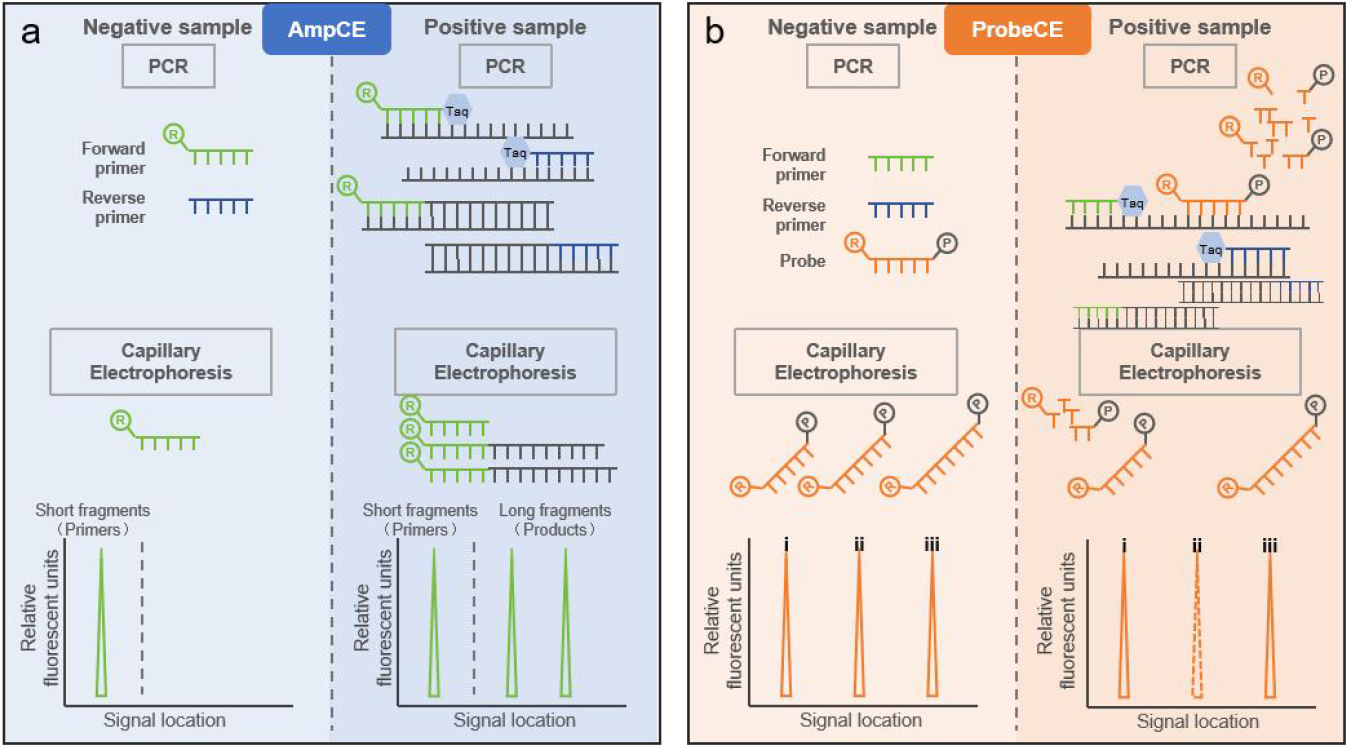
Schematic diagrams. **(a)** Schematic diagrams of conventional AmpCE. Primers with fluorescent modifications modify the amplified product through PCR reaction. AmpCE detects samples by receiving fluorescent signals from amplification products. **(b)** Schematic diagrams of ProbeCE. The probe has fluorophore at the 5th end and phosphorylation at the 3rd end. The probe is cut during the PCR process. ProbeCE detects samples through changes in probe signal intensity ®, Fluorophore. ℗, Phosphate group.

**Figure 2.**
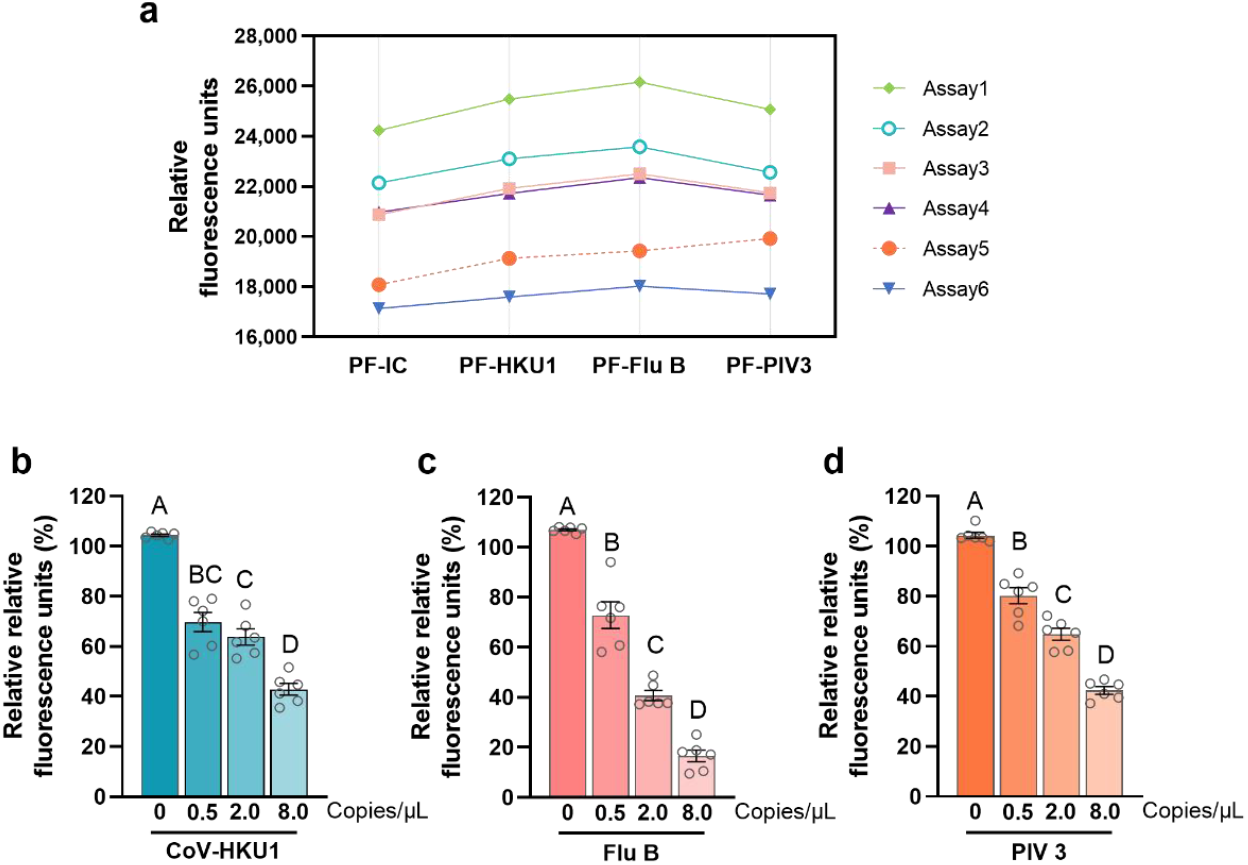
Changes in fluorescence intensity using samples with different concentrations. **(a)** Relative fluorescence units of probes in 6 replicate experiments. All test samples were water. **(b)** The ratio of signal intensity between probe PF-HKU1 and PF-IC was determined using varying concentrations of CoV-HKU1 as the template. **(c)** The ratio of signal intensity between probe PF-Flu B and PF-IC was determined using varying concentrations of Flu B as the template. **(d)** The ratio of signal intensity between probe PF-PIV3 and PF-IC was determined using varying concentrations of PIV 3 as the template. The data were presented as mean ± S.E.M., *P-value* <0.01 was considered statistically significant, *n* = 6.

**Figure 3.**
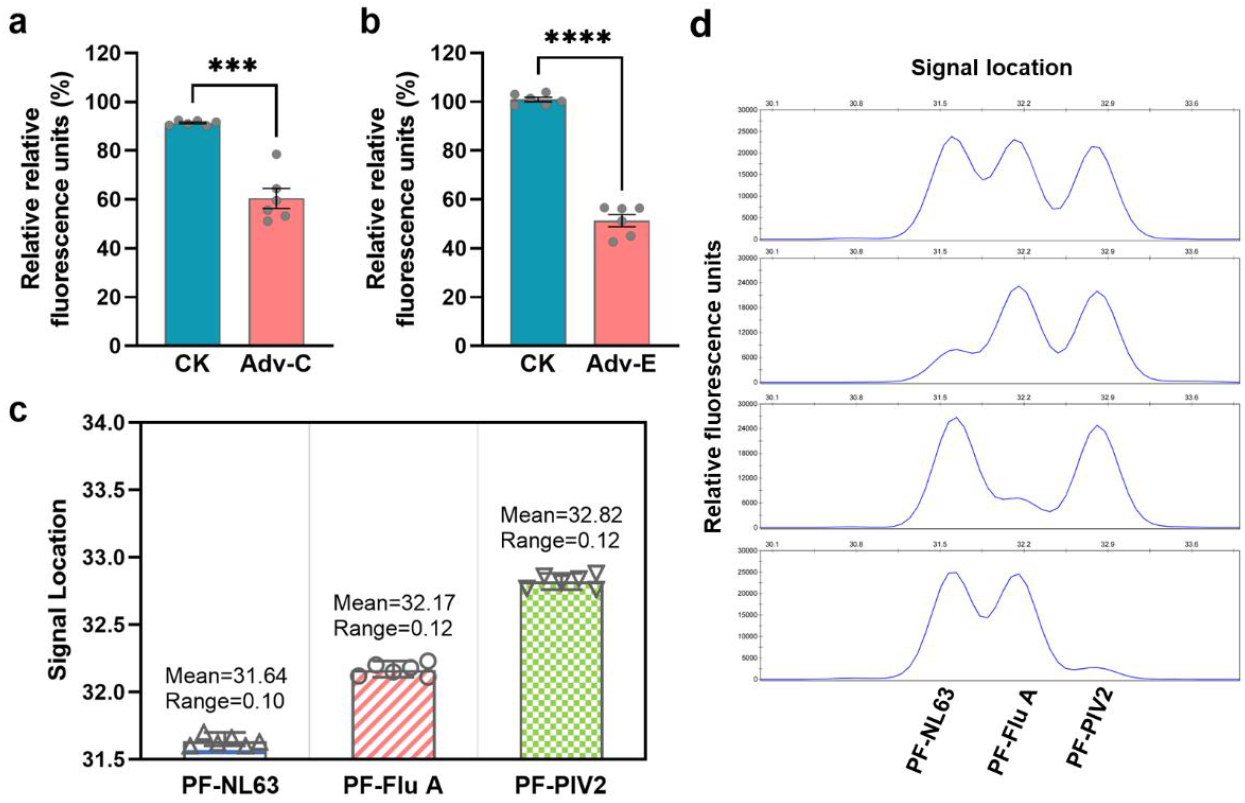
Evaluating different length probes. **(a)** The ratio of signal intensity between probe PF-Adv C and PF-IC was evaluated using water or Adv C as the template. **(b)** The ratio of signal intensity between probe PF-Adv E and PF-IC was evaluated using water or Adv E as the template. The data were presented as mean ± S.E.M., *n* = 6. **(c)** Signal location of PF-NL63, PF-Flu A, PF-PIV2. The error bar represented the Range of the Mean, *n* = 6. **(d)** Visualization of probe signals in CE.

**Figure 4.**
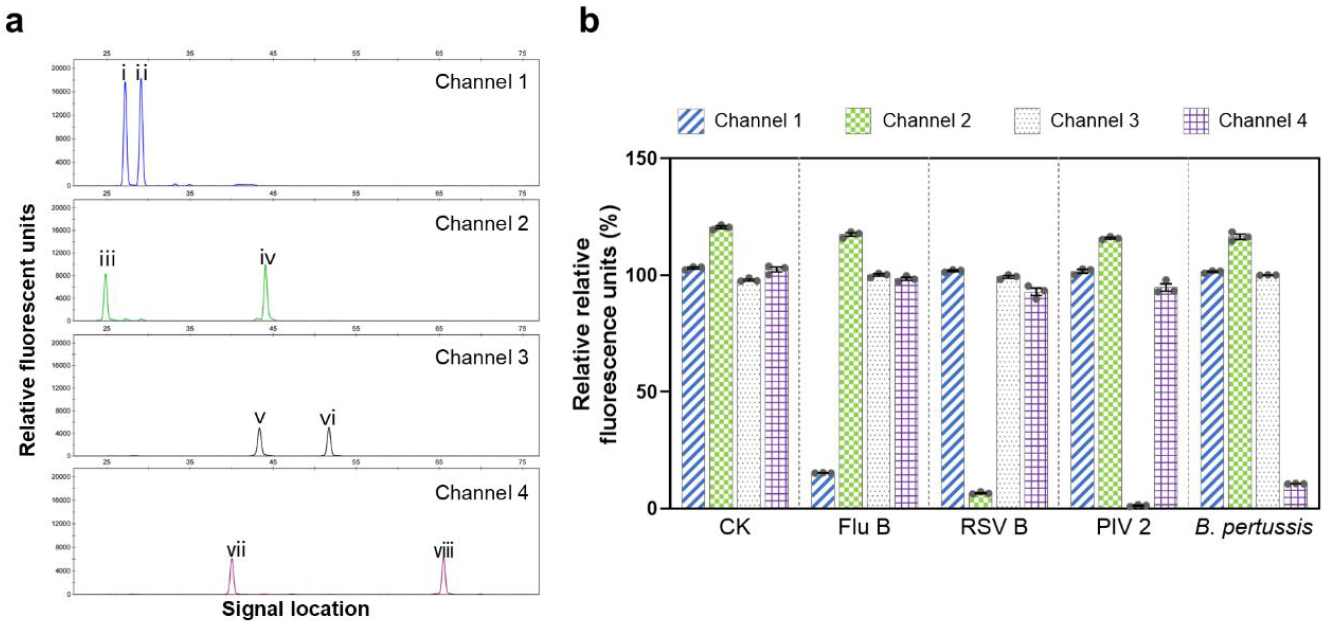
Evaluate multi-channel reception of fluorescence signals. **(a)** Visualization of multi-channel fluorescence signal in CE. Channel 1 received signals from probes modified with FAM: i was PF-HKU1, ii was PF-Flu B; Channel 2 received signals from probes modified with JOE: iii was PJ-COVID-19, iv was PJ-RSV B; Channel 3 received signals from probes modified with Atto550: v was P550-PIV2, vi was P550-PIV1; Channel 4 received signals from probes modified with Atto590: vii was P590-PIV3, viii was P590-BP. **(b)** Relative signal intensities of probes under different treatment conditions. Channel 1 displayed the ratio of PF-Flu B to PF-HKU1 signal intensity, channel 2 displayed the ratio of PJ-RSV B to PJ-COVID-19 signal intensity, channel 3 displayed the ratio of P550-PIV2 to P550-PIV1 signal intensity, channel 4 displayed the ratio of P590-BP to P590-PIV3 signal intensity. axis display of test samples: Water (CK), Flu B, RSV B, PIV 2, *B. pertussis*. The data were presented as mean ± S.E.M., *n* = 3.

The principle of ProbeCE technology relies on the 5’-3’ polymerization and exonuclease activities of Thermus aquaticus (Taq) DNA polymerase. The enzyme uses DNA as a template to synthesize new strands while removing bound DNA fragments through its exonuclease activity (15, 16). If no target nucleic acid fragment is present, DNA synthesis will not be performed and the probe will be intact. In contrast, the probe is cut during the extension phase. During the CE stage, the migration rate of the damaged probe changes, resulting in a decrease in the predicted position signal intensity.

### ProbeCE for low-concentration sample detection

The probe concentration in ProbeCE was quite low to maintain the fluorescence signal intensity within an observable range. For instance, the concentration of FAM-modified probes was 10 nM, which was 1/20th of the primer concentration.

Does such a low probe concentration affect ProbeCE’s detection of low-concentration samples? We assessed this by using a four-fold concentration gradient sample. In addition to specific primers and probes, a control probe (PF-IC), lacking corresponding primers in the reaction system, was also used in the PCR reaction system to eliminate potential errors arising from operational or instrumental sources.

While the signal intensity indicated by Relative Fluorescent Units (RFUs) from the same probe varied between experiments, the relative intensities from PF-IC, PF-HKU1, PF-Flu B, and PF-PIV3 remained unaffected **(Figure 2a)**. This indicates that the experimental conditions were sufficiently stable to allow meaningful comparisons and Relative Relative Fluorescent Units (Relative RFUs = RFUs[Tatget] / RFUs[Control]) were used to indicate changes in signal intensity. As shown in **Figure 2b-d**, Higher concentration of samples led to more significant changes in fluorescence signal intensity. In addition, when the nucleic acid concentration was 0.5 copies/μL, the change in signal intensity was still recognizable, indicating that ProbeCE can detect low-concentration samples.

### The range and spacing of the probes

In ProbeCE, probes of varying lengths were utilized. To ensure earlier binding to the nucleic acid, the melting temperature of probes must be higher than that of the primers. This study’s shortest probe was 20 nucleotides (nt) long, targeting Adv-C. While there was no restriction on probe length, this study selected 90 nt as the maximum length.

The probe had a 56-nt random sequence at the 3’ end and a 34-nt sequence at the 5’ end complementary to the template, Adv-E (**Table 1**). Both probes were cut during the extension stage when the sample contained the respective identifiable fragments, leading to reduced fluorescence signal intensity **(Figure 3a, b)**.

The resolution capability of CE influences probe placement density, as higher resolution capability allows for the placement of more probes of varying lengths within a given distance. A set of probes, differing in size by a single nucleotide, were employed to evaluate the resolution capability of CE (**Table 1**). Migration speed exhibited an inverse relationship with probe size; simultaneously, the signal location was influenced by factors including DNA base composition, secondary structure, and modifications (17), resulting in an uneven distribution. In terms of resolution, fluorescence signals could be distinguished when signal locations differed by 0.53 units **(Figure 3c, d)**.

### Multiple fluorescent channels

Since the maximum fluorescence signal intensity from the probe in ProbeCE was controllable, there is no risk of permeation due to excessive signals. It allows us to increase the number of signals by using multiple channels. We mixed probes modified with four distinct fluorophores, and fluorescence signals appeared in four independent channels **(Figure 4a)**. The fluorescence signals from the FAM-labeled probe (PF-HKU1 and PF-Flu B) were detected in channel 1, and when the sample was Flu B, the ratio of signal intensity between probe PF-HKU1 and PF-Flu B decreased. The signals from the JOE-labeled probe (PJ-COVID-19 and PJ-RSV B) were captured in channel 2, with a decrease in the ratio of signal intensity between probe PJ-RSV B and PJ-COVID-19 when the sample was RSV B. In channel 3, the signals from the Atto550-labeled probe (P550-PIV2 and P550-PIV1) were observed, and when the sample was PIV-2, the ratio of signal intensity between probe P550-PIV2 and P550-PIV1 decreased. Lastly, the fluorescence signal from the Atto590-labeled probe (P590-PIV3 and P590-BP) was detected in channel 4, with the ratio of signal intensity between probe P590-BP and P590-PIV3 decreasing when the sample was *B. pertussis* **(Figure 4)**.

## DISCUSSION

We have developed a probe and CE-based multi-gene detection method. In this method, the relative fluorescence signal intensity decreases when detectable fragments are present in the sample, while it remains unchanged in the absence of target fragments. Compared to conventional AmpCE, our method, termed ProbeCE, offers several advantages: higher specificity, simpler primer design, reduced interference signals, and increased detection capacity.

### Higher specificity

Conventional AmpCE recognizes nucleic acid fragments through forward and reverse primers (18), whereas ProbeCE utilizes forward primer, reverse primer, and probe for recognition. This significantly reduces the probability of misidentification.

### Simplified primer design

In conventional AmpCE, sample identification is based on the difference in the amplification product length, requiring precise control over the amplification window. In contrast, ProbeCE has no such limitations.

Although ProbeCE requires additional probes, it does not involve fluorescence quenching with no length restriction, making it less complex than TaqMan qPCR, which relies on fluorescence resonance energy transfer (FRET) (2).

### Reduced interference signals

In conventional AmpCE, the fluorescence signal is generated from residual primers, specific amplification products, and non-specific amplification products. Distinguishing between non-specific amplification signals and specific target product signals can be difficult, particularly in a multiplex PCR system that employs multiple primers, which increases the likelihood of non-specific amplification. Additionally, saturation of the fluorescence signal can result in peak-dragging artifacts (19). All of this makes conventional methods susceptible to interference. In contrast, ProbeCE focuses on the fluorescence signal from the probe, which maintains a stable signal position and can only be cut. While interference signals from the damaged part may occur, these can be predicted and managed since the probe sequence is known.

### Increased detection capacity

CE supports multicolor fluorescence detection (8). However, Conventional AmpCE cannot apply this for non-quantitative samples due to signal saturation, which may interfere with other channels. Moreover, the length of the amplified product is unstable due to base mutations, deletions, and insertions, requiring a larger signal spacing for gene identification. In comparison, the fluorescence signals from the probe are more stable and a 0.53-unit spacing is sufficient for reliable resolution (**Figure 3**).

In detecting known genes, ProbeCE offers advantages in usability and cost over current medium to high-throughput gene identification methods.

1. **More targeted:** Using universal/specific primers and specific probes, ProbeCE focuses research data on a defined area of interest.
2. **User-friendly:** Experiments are performed with fewer steps and results are presented visually. In addition, the PCR amplifier and CE equipment required for the experiment are relatively common.
3. **Cost-effective:** Although ProbeCE requires consumables for CE, its overall costs are considerably lower than those of microfluidic systems or other methods. In addition, the simple detection process greatly reduces labor costs, making ProbeCE a more economical option.

However, ProbeCE does have its limitations: 1) This method can compare the copy numbers of nucleic acids within a certain range **(Figure 2)**, but it does not support sample quantification. 2) ProbeCE can only distinguish known sequence differences and cannot discover new types of mutations. Nevertheless, we believe this method has great potential for widespread application in the future.

## DATA AVAILABILITY

The data is available upon request from the authors.

## AUTHOR CONTRIBUTIONS

Yangqing Gong: Conceptualization, Formal analysis, Methodology, Validation, Writing—original draft. Yong Wu: Resources. Yong Luo: Supervision, Writing—review & editing. Xinjiang Lu: Supervision, Writing—review & editing. Miaomiao Niu: Resources.

## ACKNOWLEDGEMENTS

We thank Haowei Zhu for kind gift of standard strains.

