## Supplementary Table 1 for "Novel Probe Design for Multi-Gene Detection Enabled by Cleavage and Capillary Electrophoresis"

| **Primer** | **Sequence (5’-3’)** |
| --- | --- |
| CoV-HKU1 | Forward: GAAGCATGGTATGAATTTCGTG |
|  | Reverse: ACCTGCACAATTACAGCCAA |
| Flu B | Forward: GGAGGAAGTAAACACTCAGAAAGA |
|  | Reverse: CCACTCTGGTCATATGCATTCAATCT |
| PIV 3 | Forward: CTTGAACCATTGCAGTCACC |
|  | Reverse: CTCGTCTTGAAGGAAGTACAATCTA |
| ADV-C | Forward: CTACAAGCGCGTGTATGATGAG |
|  | Reverse: CATGTCCTTATGCCGCTTTC |
| ADV-E | Forward: GACAGAACCCGGTATTTCAG |
|  | Reverse: AGTTTGGCAATTCATCCTCCAC |
| CoV-NL63 | Forward: CAACACGTCCATCATTAACATACC |
|  | Reverse: GCTTGTACTAGGTTGGAAGGT |
| Flu A | Forward: TCTTCTAACCGAGGTCGAAACGTA |
|  | Reverse: GTCTTGTCTTTAGCCATTCCA |
| PIV 2 | Forward: AGCCACAACACTTAAATTCACTC |
|  | Reverse: GATGAGCTAGTGTGCAAATGG |
| RSV-B | Forward: TCAGTGTTACAAAGGCTGACT |
|  | Reverse: ATATGTTGTACAGCTACCTATC |
| COVID-19 | Forward: CAGGCAAGTTAAGGTTAGATAGCA |
|  | Reverse: TGAAGATTAATGCGGCTTGTAG |
| PIV 1 | Forward: GCAGTGACATCCAGAGTTCC |
|  | Reverse: CTTGTTAAAGCTTTGAGGCACT |
| PIV 4 | Forward: CTACTTCAGCAAATTTGACCCA |
|  | Reverse: GCTTAATAGGCAATGCATTCCA |
| BP | Forward: GGAGGCGAGAATAAGACAATCTA |
|  | Reverse: GCGTCTATTACATTTCCGGTCA |

**Supplementary Table 1. Sequence of Primers used in ProbeCE**
