## Supplementary Table 2 for "Novel Probe Design for Multi-Gene Detection Enabled by Cleavage and Capillary Electrophoresis"

**Supplementary Table 2. Sequence of DNA ladder used in CE**

| **Name** | **Sequence (5'-3')** | **Size (nt)** |
| --- | --- | --- |
| Size 20 | Alexa fluor 633-ATAAGACTCGGCGGTTACTC-P | 20 |
| Size 40 | Alexa fluor 633-AGTGTAAGACTCGGCGGTTTGGTACTTCAC ACTACACACG-P | 40 |
| Size 70 | Alexa fluor 633-TCCATCAGGACGGGGAATAACTATCTCCAA CGTACATTGGCACTGATTTAGATGGTGACGACCTGTGCCG-P | 70 |
